# Evaluating zero-shot prediction of protein design success by AlphaFold, ESMFold, and ProteinMPNN

**DOI:** 10.1101/2025.07.29.667290

**Authors:** Mario Garcia, Sugyan M. Dixit, Gabriel J. Rocklin

## Abstract

*De novo* protein design has enabled the creation of proteins with diverse functionalities that are not found in nature. Despite recent advances, experimental success rates remain inconsistent and context-dependent, posing a bottleneck for broader applications of *de novo* design. To overcome this, structure and sequence prediction models have been applied to assess design quality prior to experimental testing to save time and resources. In this study, we aimed to determine the extent to which AlphaFold2, Protein MPNN, and ESMFold can discriminate between experimentally successful and unsuccessful designs. For this, we curated a benchmark dataset of 614 experimentally characterized *de novo* designed monomers from 11 different design studies between 2012 and 2021. All predictive models demonstrated moderate ability to discriminate experimental successes (expressed, soluble, monomeric, and fold into the intended design structure) from failures, with many failed designs having better confidence metrics than successful designs. Among all computational models evaluated, ESMFold average pLDDT yielded the best individual performance at distinguishing between successful and unsuccessful designs. A logistic regression model combining all confidence metrics provided only modest improvement over ESMFold pLDDT alone. Overall, these results show that these models can serve as an initial filtering strategy prior to experimental validation; however, their utility at accurately predicting experimental successful designs remains limited without task-specific training.

## Introduction

*De novo* protein design permits the exploration of protein sequence space beyond the constraints imposed by natural evolution (1). This approach allows for the design of novel protein folds with precise control over the designed structure to achieve specific functions, representing promise for developing novel proteins with applications in medicine and biotechnology (2). Despite recent breakthroughs in AI-guided structure prediction and design strategies, experimental success rates of designs remain inconsistent with rates varying based on the complexity of the designed topology and design objective (3). For example, α-helical protein designs tend to have higher experimental success rates compared to designs rich in β-sheet content or designs attempting to incorporate functionality (3, 4). These inconsistencies emphasize the need for improved computational models to better predict which designs will likely succeed experimentally.

Protein design can fail at many experimental stages, including through poor expression, low solubility, aggregation, folding to an undesired or non-functional conformation, or low overall folding stability. Implementing computational strategies to filter out bad designs prior to characterization can reduce the experimental resources and cost associated with validating designs (5, 6).

Over the last several years, deep learning models have increasingly been applied for this purpose. Structure-based prediction models such as AlphaFold, inverse folding prediction models such as Protein MPNN, and protein language models such as ESMFold have been integrated in design strategies as final validation steps to remove poor designs from experimental characterization (7-12). Although these computational models have shown great capabilities at sequence and structure prediction, they were not trained to predict protein biophysical properties such as expression, aggregation, or stability. As a result, many *de novo* designed proteins achieve highly confident model metrics even when these designs fail to express or fold under typical experimental conditions (13-15). In one large-scale test on *de novo* designed αββα miniproteins, AlphaFold pLDDT metric showed minimal discrimination between experimentally stable and unstable designs (16). These examples highlight the importance of assessing the performance of sequence and structure prediction models when tasked to discriminate between experimental successes and failures in *de novo* design. Furthermore, design and prediction are separate challenges. Even if a protein design method arose with near-perfect success at a certain design task, it would remain important to accurately predict whether diverse sequences generated by other approaches could also achieve the design objective.

In this study, we aim to quantify the extent to which existing deep learning models developed for structure and sequence prediction can distinguish between experimentally successful and unsuccessful *de novo* protein designs. To accomplish this, we compiled a dataset of 614 experimentally characterized *de novo* designed proteins covering many different designed topologies from the past decade. All proteins were designed using Rosetta-based methods to fold into soluble monomers without deep learning, indicating that these designs all had strongly favorable energies according to a classical protein design energy function (17). Since these models have never been trained on experimental outcomes such as protein expression, solubility, or folding stability, asking them to predict design success is a “zero-shot” evaluation of these models. This analysis provides a baseline for their performance in predicting design success. We also use our benchmark dataset containing many unique folds to examine the influence of protein topology on model confidence and accuracy.

## Methods

### Dataset Construction

The design studies selected for this analysis focused on the *de novo* design of monomeric proteins (18-28). These studies did not include AlphaFold2, Protein MPNN, or ESMFold during the design process. All monomeric designs in the dataset were experimentally characterized using expression in E. coli, protein purification, size exclusion chromatography combined with multi-angle light scattering (SEC-MALS), and secondary structure assessment by circular dichroism (CD). We defined a design to be experimentally successful if it was expressed, soluble, monomeric, and had a CD spectrum consistent with the intended design fold as reported in each study (Fig. 1A). Protein designs that did not report experimental success or designs needing to be stabilized as dimers were excluded from the final analysis.

**Figure 1.**
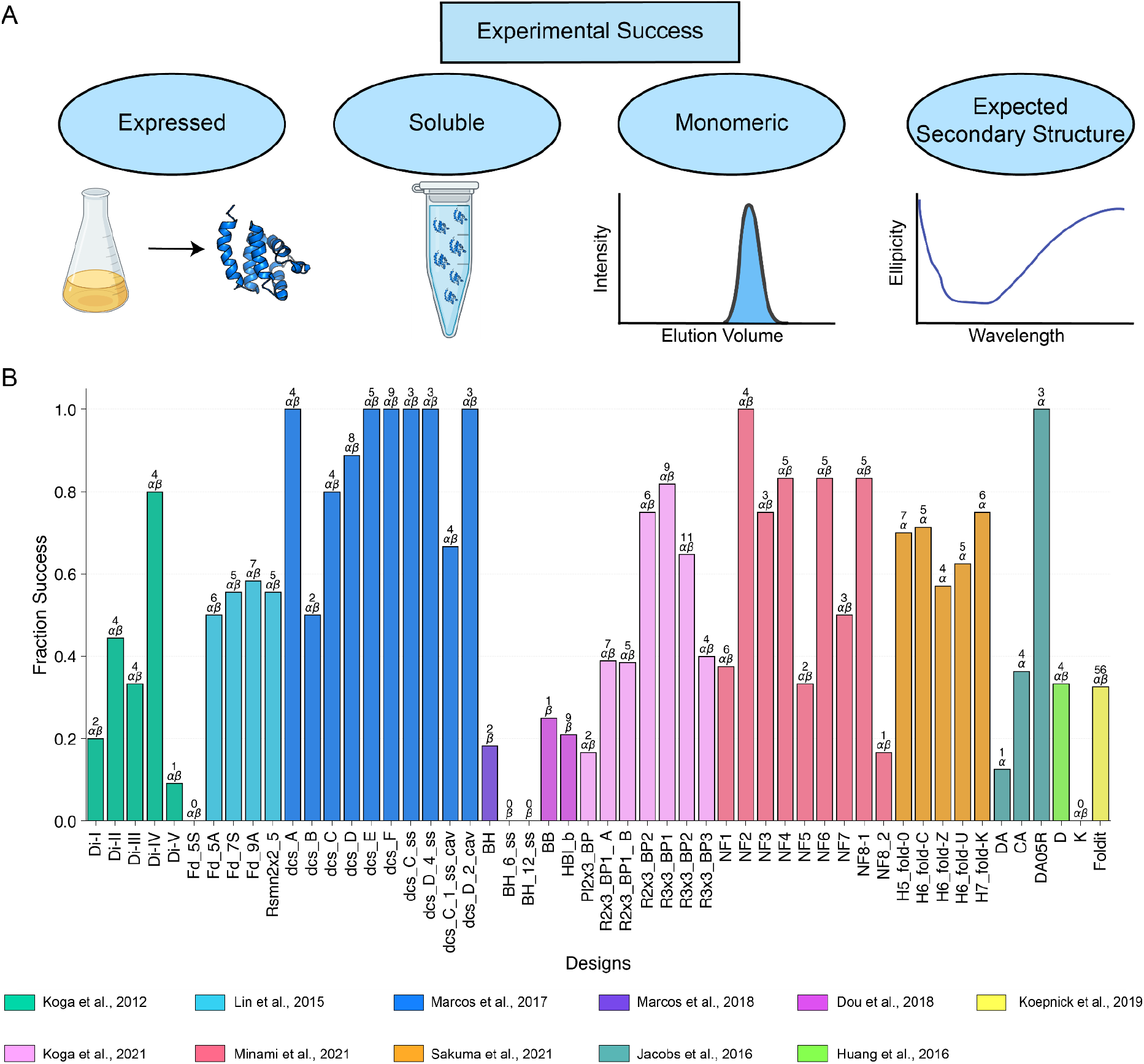
Experimental success rates vary depending on design objective and topology. **A)** Schematic depiction of criteria used to define experimentally successful de novo designs in this analysis. A design was considered successful if it was expressed, soluble, monomeric, and folded into the intended secondary structure **B)** Fraction of experimentally successful designs for each design topology. Design topologies consisted of all α-helical proteins (α), all β-sheet proteins (β), or mixed α/β proteins (α/β). The number of successful designs within each topology is reported above each bar. Bars for each design topology are colored according to the reported article.

### AlphaFold Modeling

AlphaFold2 (AlphaFold 2.1.1 version) was used to generate predicted structures for all monomeric *de novo* designs in the dataset (7). For each sequence, five predicted protein structures were generated using the AlphaFold monomer model. The confidence for each structure was determined by the pLDDT metric. The structure with the highest average pLDDT value was selected as the best-ranked AlphaFold predicted structure. AlphaFold structures were generated using both the default protocol of 3 recycles and an extended protocol with 25 recycle steps (7). In addition, AlphaFold PTM model was used to compute the average PAE of each protein design. The average AlphaFold pLDDT and PAE of each design were used as confidence metrics to assess model performance when predicting experimental success. A higher AlphaFold pLDDT and a lower PAE indicate better model predictive performance.

### ProteinMPNN Modeling

We used the highest-ranked AlphaFold predicted structures with the default recycle parameter as the input backbone for Protein MPNN (9). The computed Protein MPNN score for the original sequence was used to assess the model’s performance in predicting experimental success, with a lower Protein MPNN score indicating better performance.

### ESMFold Modeling

To assess the ability of protein language models at predicting experimental success, ESMFold was applied to all *de novo* designed proteins in the dataset (8). ESMFold (esmfold_v1) was used to predict the structures for all designs. ESMFold performance was assessed using the average pLDDT of the predicted structures. A higher average pLDDT indicates better model performance for ESMFold.

### Logistic Regression Modeling

Regression models and ROC curves were computed using scikit-learn (29).

## Results

### Roughly half of the designs across 11 studies met all experimental success criteria

We compiled 11 *de novo* protein design studies that collectively included 614 individual monomeric *de novo* designs. Each study typically focused on several different design “topologies” specifying the lengths and connectivity of secondary structural elements in each design. Overall, 269/614 (43%) were experimentally successful (soluble, monomeric, and with the correct secondary structure). Success rates varied from study to study and topology to topology (Fig. 1B). For example, whereas most (88%) of the curved β-sheet proteins in Marcos et al. (22) met all experimental success criteria, a much smaller fraction (21%) of the β-barrel designs from Dou et al. (24) were experimentally successful. Many of the lowest success rates came from efforts to design β-sheet rich proteins, such as β-barrel and jellyroll designs (28). Most failed designs were attributed to insolubility and aggregation (65% of failures) (Fig. S1 & S2), although many soluble, monomeric designs failed to show the intended secondary structure via circular dichroism (59/345). Figure 2 displays examples of soluble designs predicted to be well-folded with high confidence by AlphaFold but failed to adopt the designed secondary structure. These designs indicate that soluble, monomeric expression is not sufficient to establish proper folding.

**Figure 2.**
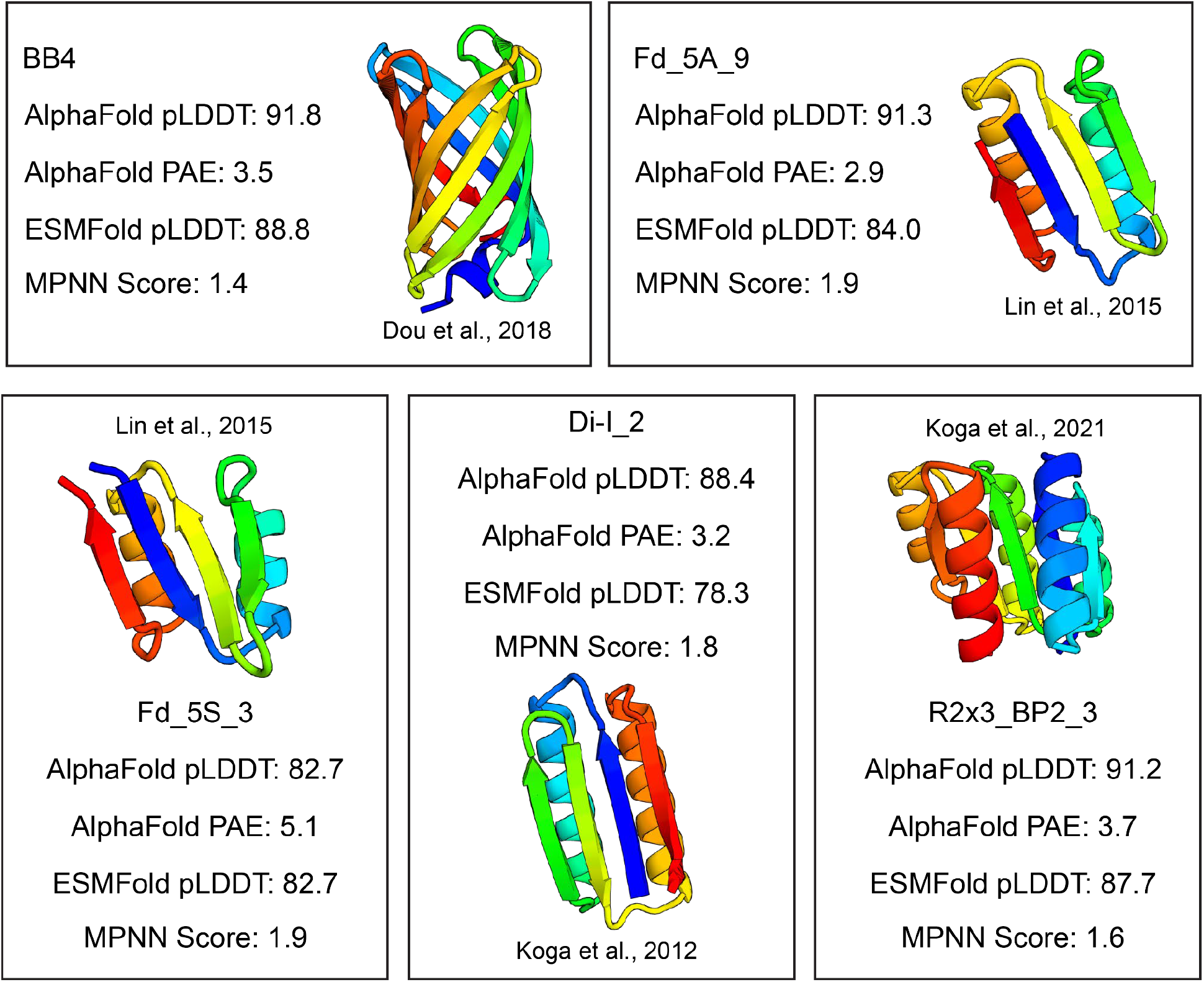
Soluble designs that failed to fold into intended secondary structures despite high AlphaFold confidence. High confidence (pLDDT > 82) AlphaFold predicted structure of designs that were experimentally soluble but failed to fold into intended secondary structures, assessed by circular dichroism.

### Structure prediction confidence metrics moderately discriminate between good and bad designs

Overall, experimentally successful designs had better confidence metric distributions compared to unsuccessful designs across all computational models (Fig. 3A). Still, discrimination was far from perfect, with ROC AUC scores ranging from 0.66-0.73 for the different confidence metrics (Fig. 3B). ESMFold’s average pLDDT had the highest AUC out of the different confidence metrics (0.73±0.04, mean±std from bootstrapping) whereas MPNN score had the lowest AUC (0.66±0.05). AlphaFold pLDDT values generated with different numbers of recycles were highly correlated (Spearman = 0.94). A total of 31/614 designs showed an increase in pLDDT > 5 using 25 recycles, with three designs increasing their pLDDT > 15. These 31 designs were primarily FoldIt designs (23/31) from Koepnick et al. (Fig. S3). The different confidence metrics captured different information, with Spearman correlations between metrics ranging from 0.52-0.83 (not including the two AlphaFold protocols, Fig. 3C). Still, despite differences between these methods, we found that combining these metrics in a logistic regression model (ROC AUC of 0.74 ± 0.04) showed minimal improvement over the best individual method (ESMFold: ROC AUC of 0.73 ± 0.04) (Fig. 4).

**Figure 3.**
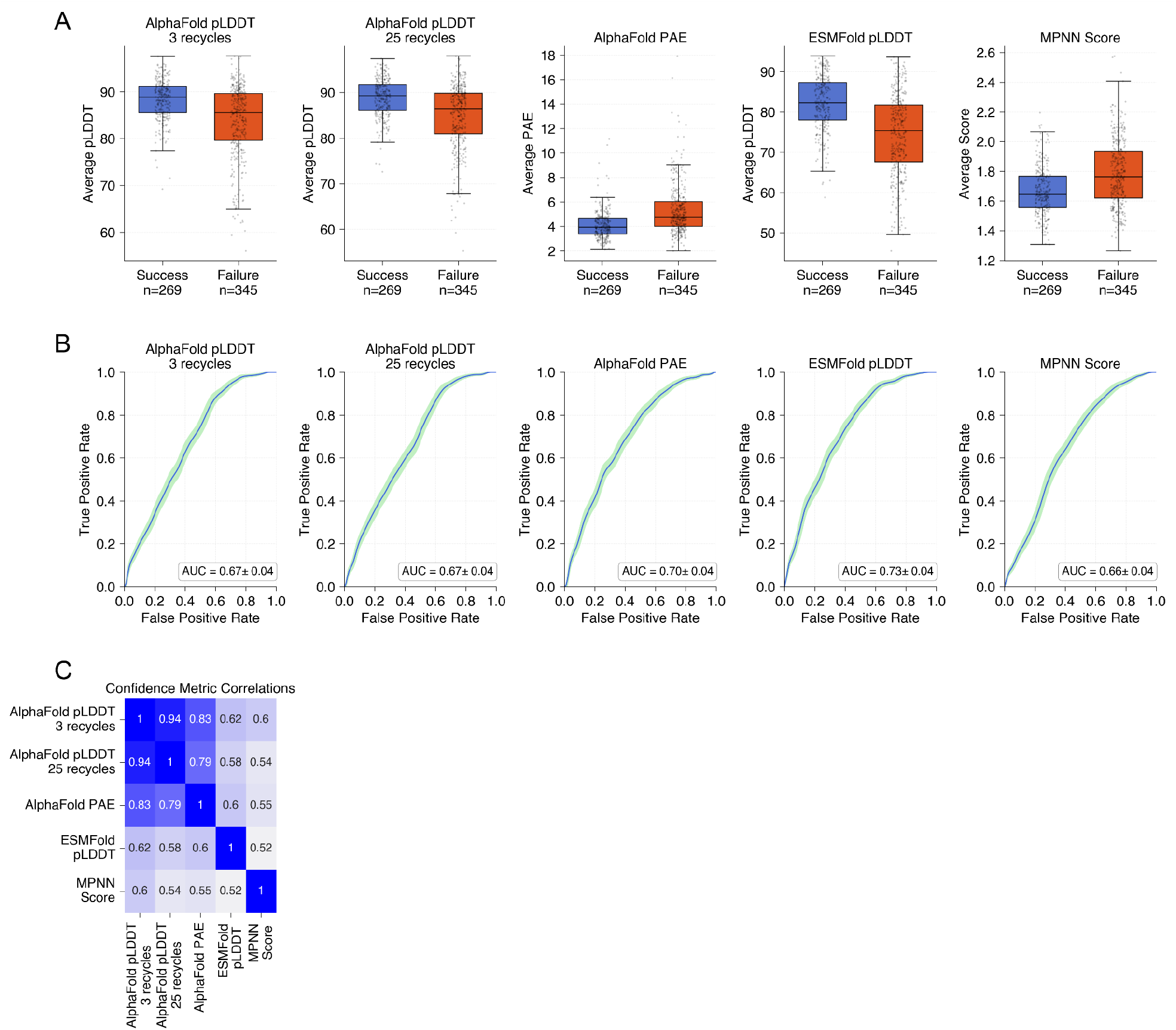
ESMFold pLDDT provides the largest discrimination at distinguishing between experimental successful and unsuccessful designs. **A)** Boxplots displaying distributions of confidence metrics between experimentally successful (blue) and unsuccessful (orange) designs with successful designs displaying better confidence metrics on average. **B)** ROC curves for each confidence metric displaying the average AUC values. The shaded region represents the 95% confidence interval computed using bootstrapping. **C)** Spearman correlation matrix showing absolute correlation between different confidence metrics.

**Figure 4.**
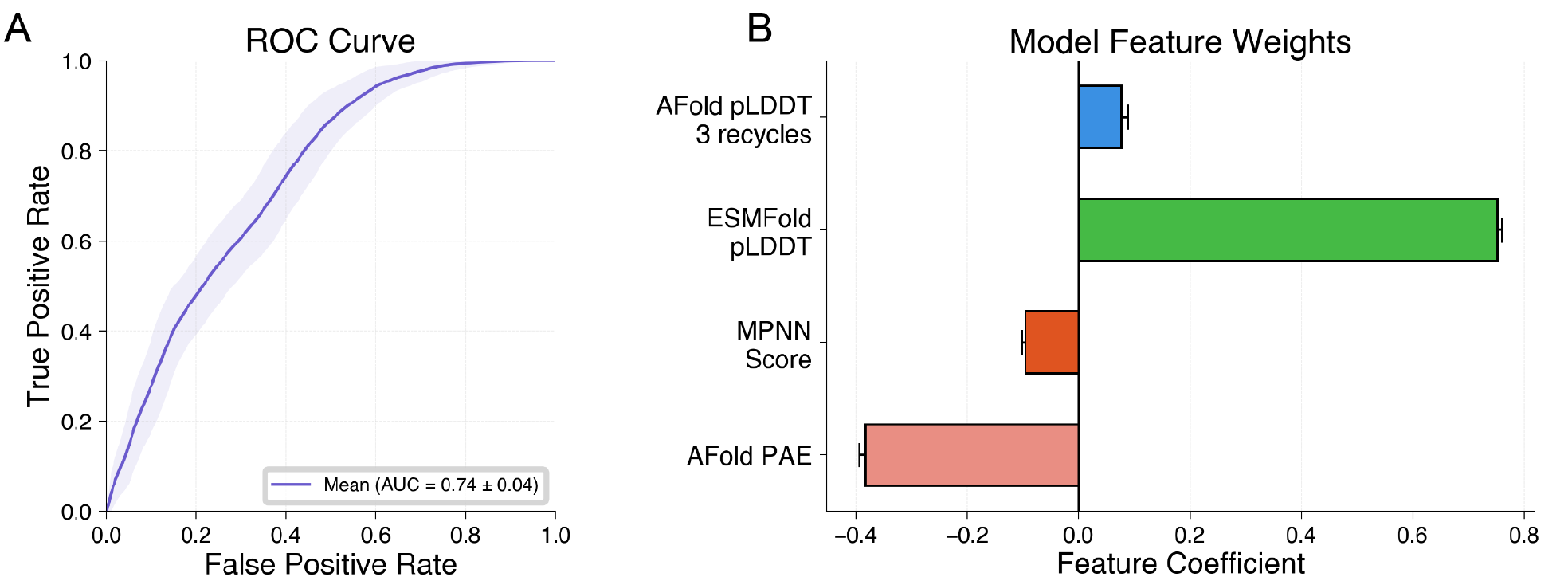
Combining confidence metrics into logistic regression model leads to minor improvement in average AUC. **A)** ROC Curve for logistic regression model trained on all confidence metrics to predict experimental success. The ROC Curve displays the mean (dark blue) and 95% confidence interval (shaded) generated through bootstrapping. **B)** Feature weights from the logistic regression model indicating the relative contribution of each metric to the model. ESMFold pLDDT showed the highest feature weight.

### Confidence metrics were not generalizable across design topologies, preventing a reliable threshold from being established for experimental success

We then analyzed confidence metrics across different design topologies, focusing initially on AlphaFold average pLDDT values (Fig. 5, Fig. S4-S6). Importantly, we observed that the distribution of AlphaFold pLDDT values varied across different design topologies (Fig. 5A). This topology-dependence indicates that there is no clear “threshold” in AlphaFold pLDDT (or ESMFold pLDDT, Fig. S5) for experimental success. Failed designs from “easily predictable” topologies on the left of Fig. 5 often had higher pLDDT scores than successful designs from “difficult to predict” topologies on the right on Fig. 5. Although AlphaFold (and ESMFold) pLDDT scores remain useful ranking metrics within an individual topology, this illustrates the difficulty of comparing designs between topologies and using pLDDT as a proxy for experimental success.

**Figure 5.**
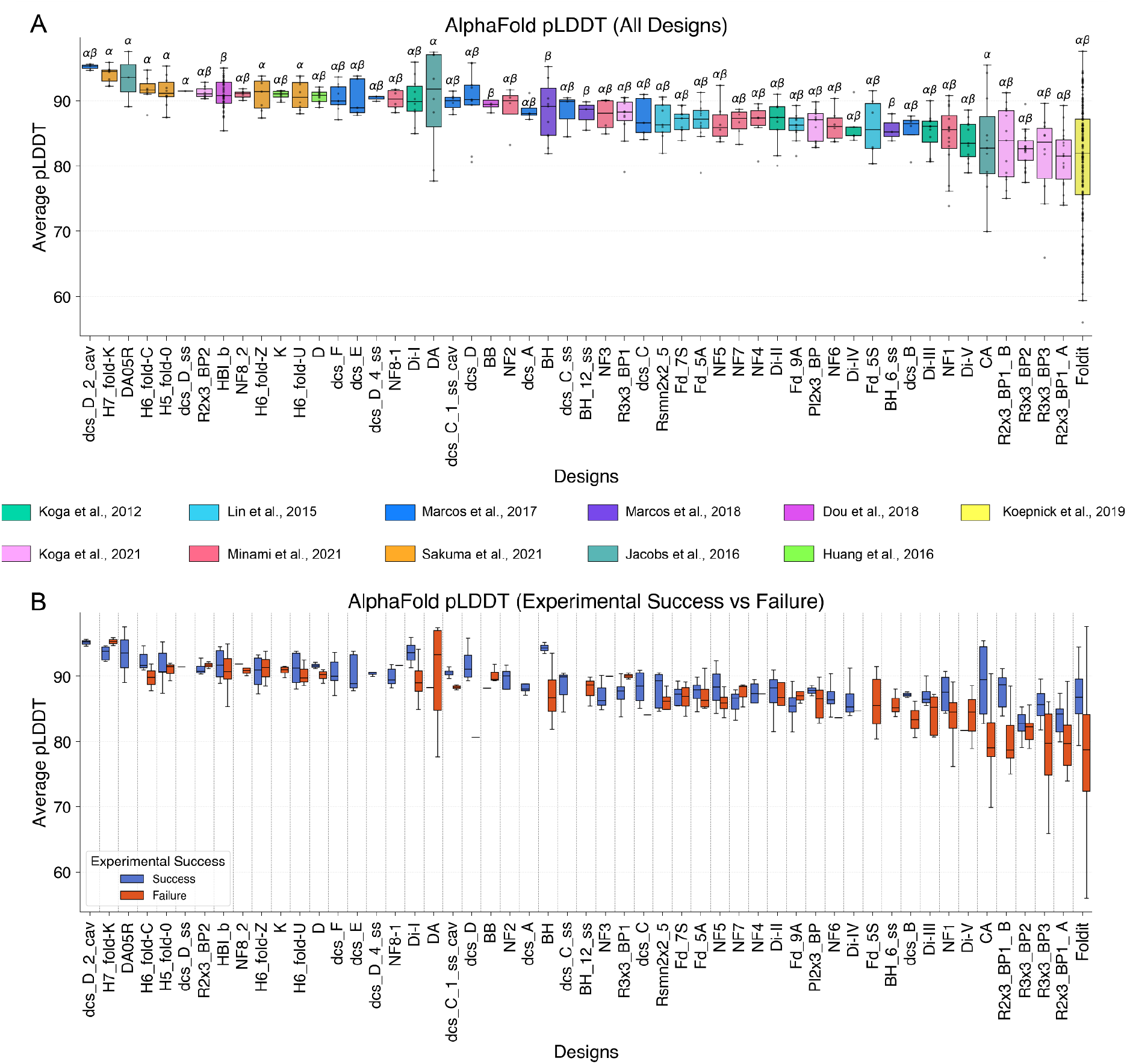
AlphaFold average pLDDT distributions vary across different design topologies. **A)** Distribution of AlphaFold average pLDDTs for designs included in this study. **B)** Distribution of AlphaFold average pLDDTs comparing experimentally successful designs (blue) and unsuccessful designs (orange) for each design topology.

## Discussion

In this study, we found that sequence and structure prediction models offer a moderate ability to distinguish between experimentally successful and unsuccessful *de novo* designs, with ESMFold providing the highest discrimination. Integrating confidence metrics from sequence and structure prediction models into a logistic regression model resulted in marginal improvements compared to ESMFold pLDDT alone. Notably, many unsuccessful designs received better confidence metrics than successful designs across all models, and confidence metrics were correlated with the design topology.

A caveat to this analysis is that many of the successful designs with crystal structures were likely implemented in AlphaFold’s training set. This would potentially bias the models to more confidently predict these structures and artificially inflate discriminatory power. However, AlphaFold’s confidence metrics struggled to consistently distinguish experimentally successful designs from failures. Examining the predicted structures of these models can potentially overcome these inconsistencies. For this analysis, we did not have access to the intended designed backbones; however, RMSD calculations between predicted backbone structures and the intended backbone designs can be implemented in future work to identify large deviations from the intended backbone design to further filter poor designs (6). Moreover, computational models are continuously being developed to predict experimental properties such as solubility, conformational dynamics, and stability (30-33). Incorporating these models into design strategies can further improve the experimental success rates. As large-scale experimental datasets become available, we anticipate that training and fine-tuning these models on this data will help improve the inconsistencies in experimental success rates of *de novo* designs. This work provides a baseline analysis of experimental success rates for *de novo* designs.

## Supporting information

denovodesigns

---

**Figure S1.**
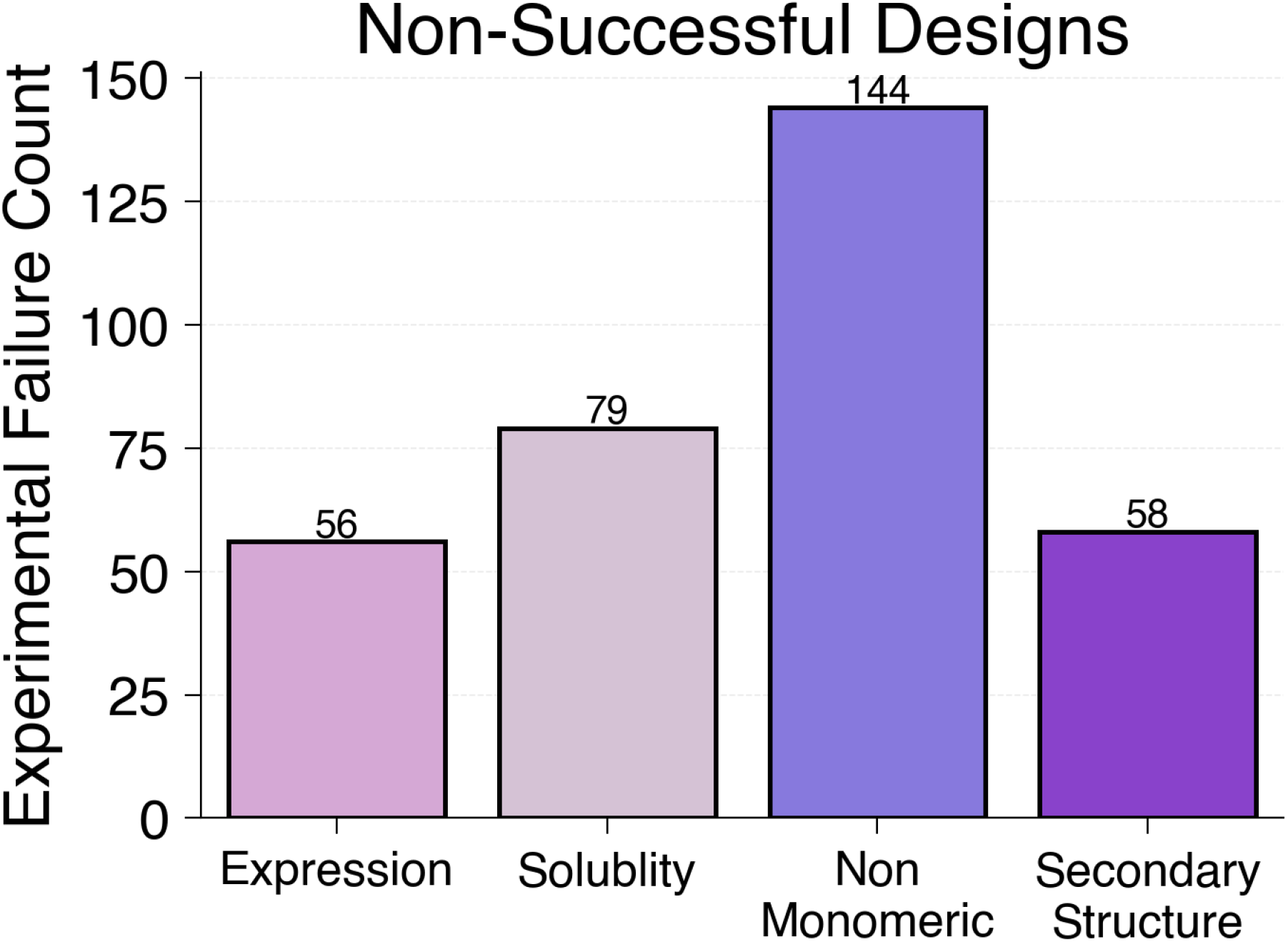
The majority of de novo designs failed experimentally due to not being monomeric. Counts of experimentally unsuccessful designs for each experimental criteria used to define design experimental success.

**Figure S2.**
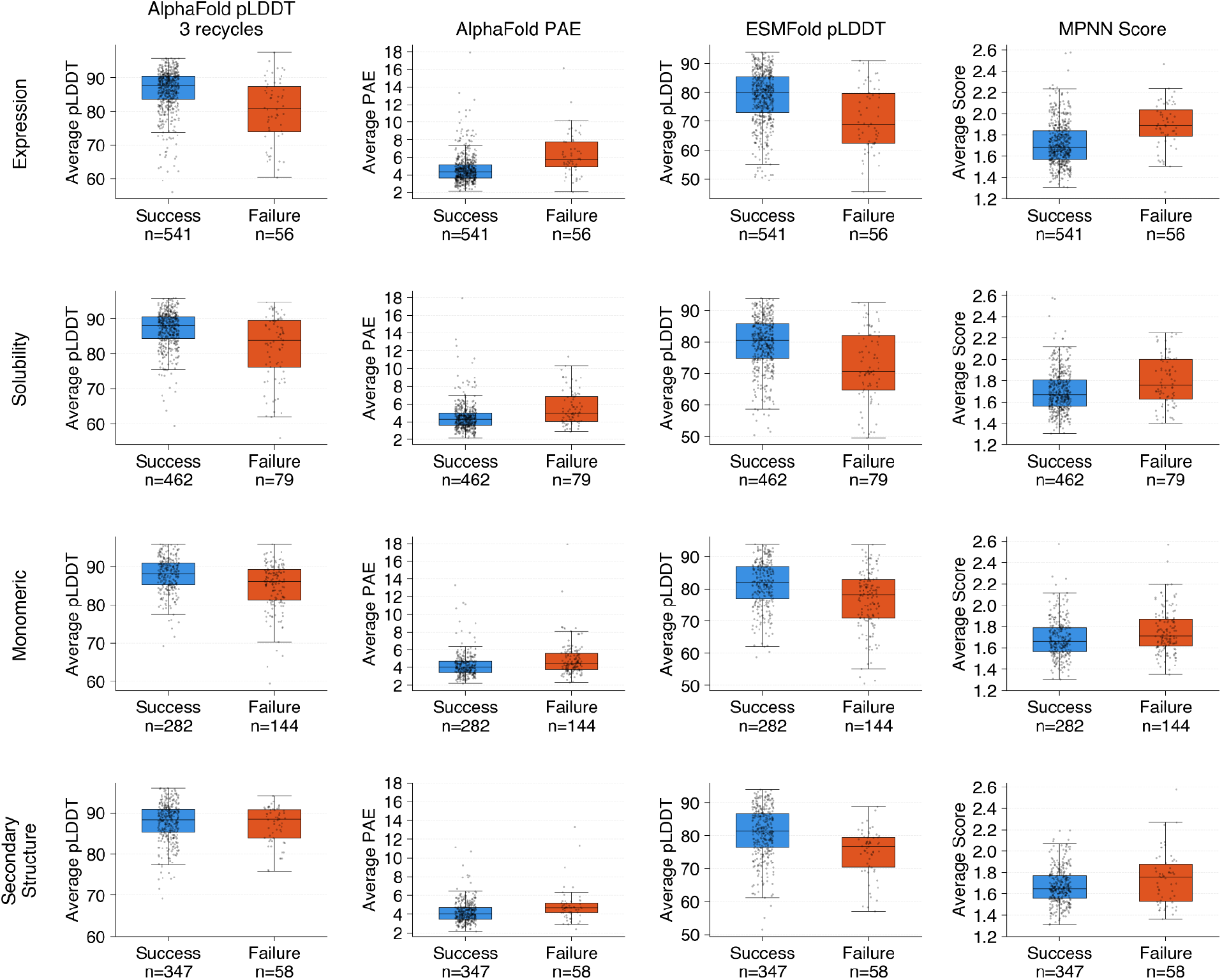
Distribution of confidence metrics for experimental outcomes of designs. Distribution of confidence metrics comparing successful (blue) and unsuccessful designs (orange) across different experimental outcomes: expression, solubility, monomeric state, and secondary structure.

**Figure S3.**
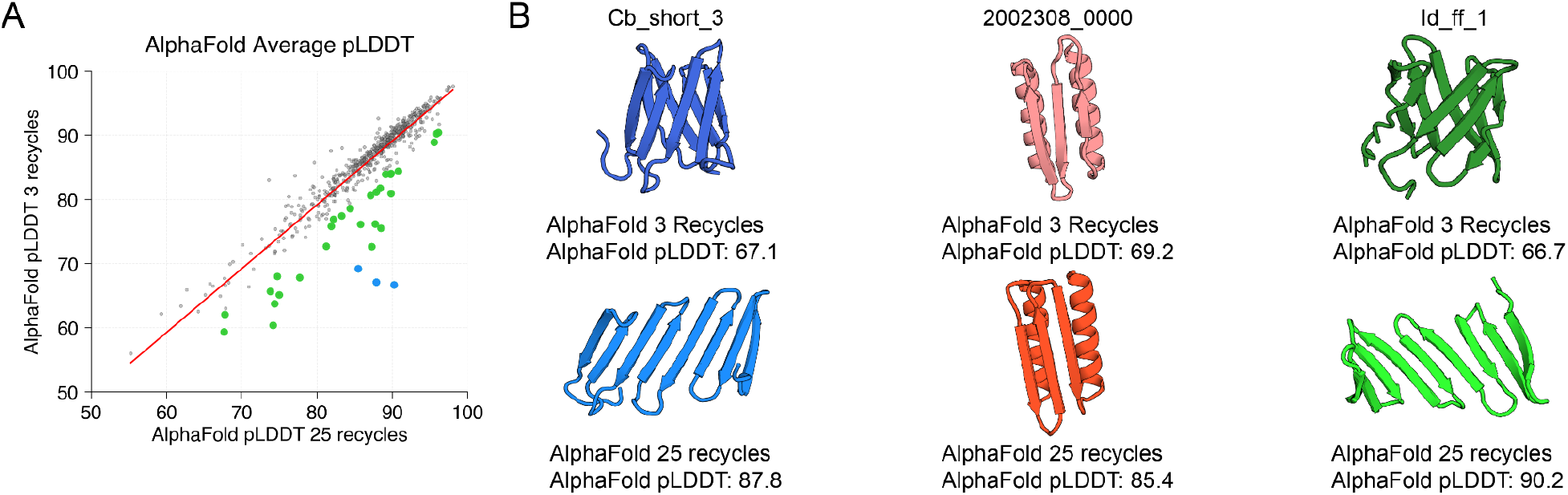
Average AlphaFold pLDDT values are highly correlated with different recycle parameters. **A)** Scatterplot capturing the relationship between AlphaFold average pLDDT for de novo designs computed using different recycle parameters. Green highlighted points indicate designs that increased their average pLDDT by >5 after increasing the recycles, and blue highlighted points are designs that increased their average pLDDT by >15 after increasing recycles. **B)** AlphaFold predicted structures of designs that increased their average pLDDT >15 with increased recycles.

**Figure S4.**
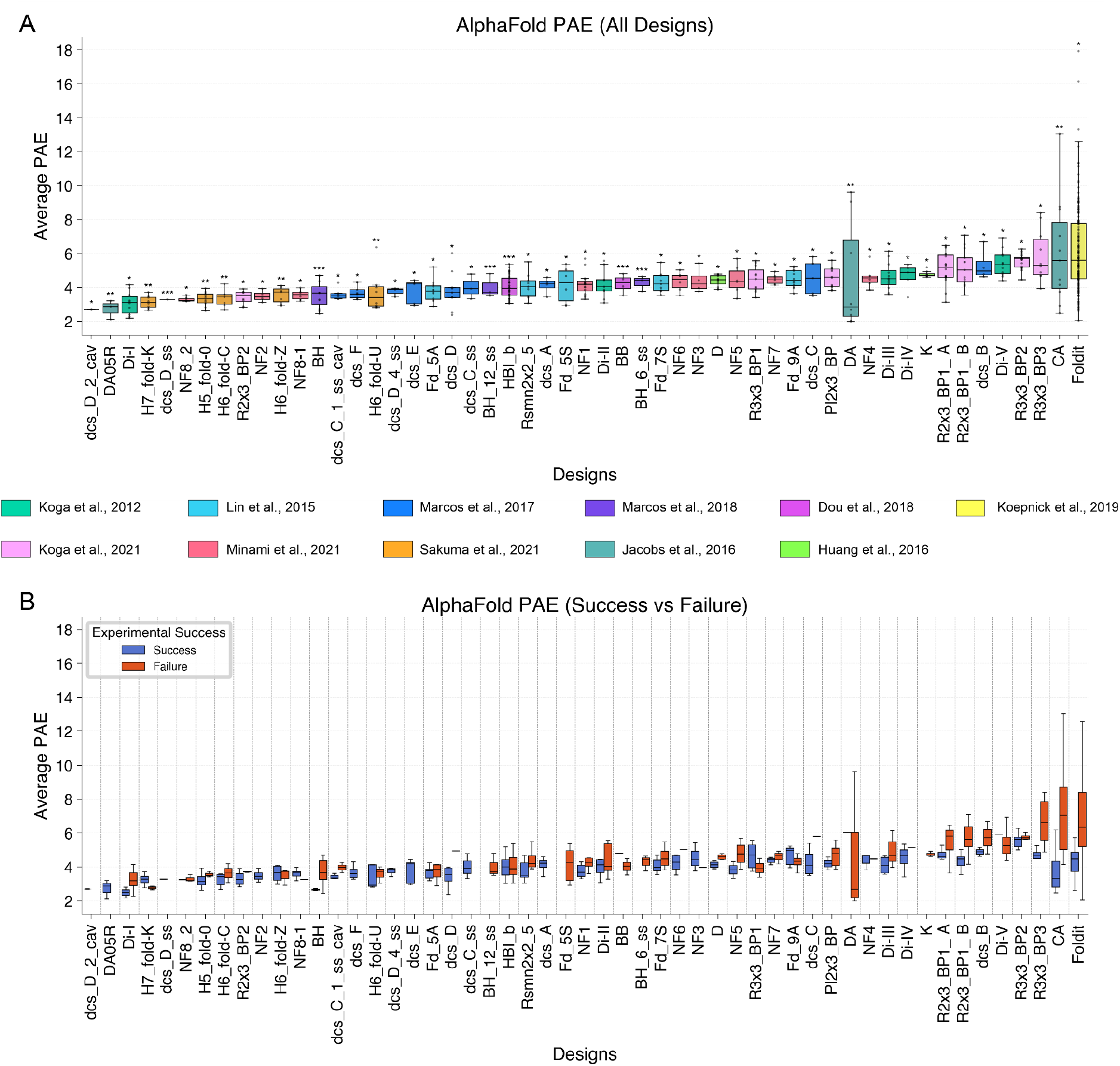
AlphaFold average PAE distributions for designed topologies. **A)** Distributions of AlphaFold average PAE for designs included in this study. **B)** Distribution of AlphaFold average PAE for experimentally successful designs (blue) and unsuccessful designs (orange) for each design topology.

**Figure S5.**
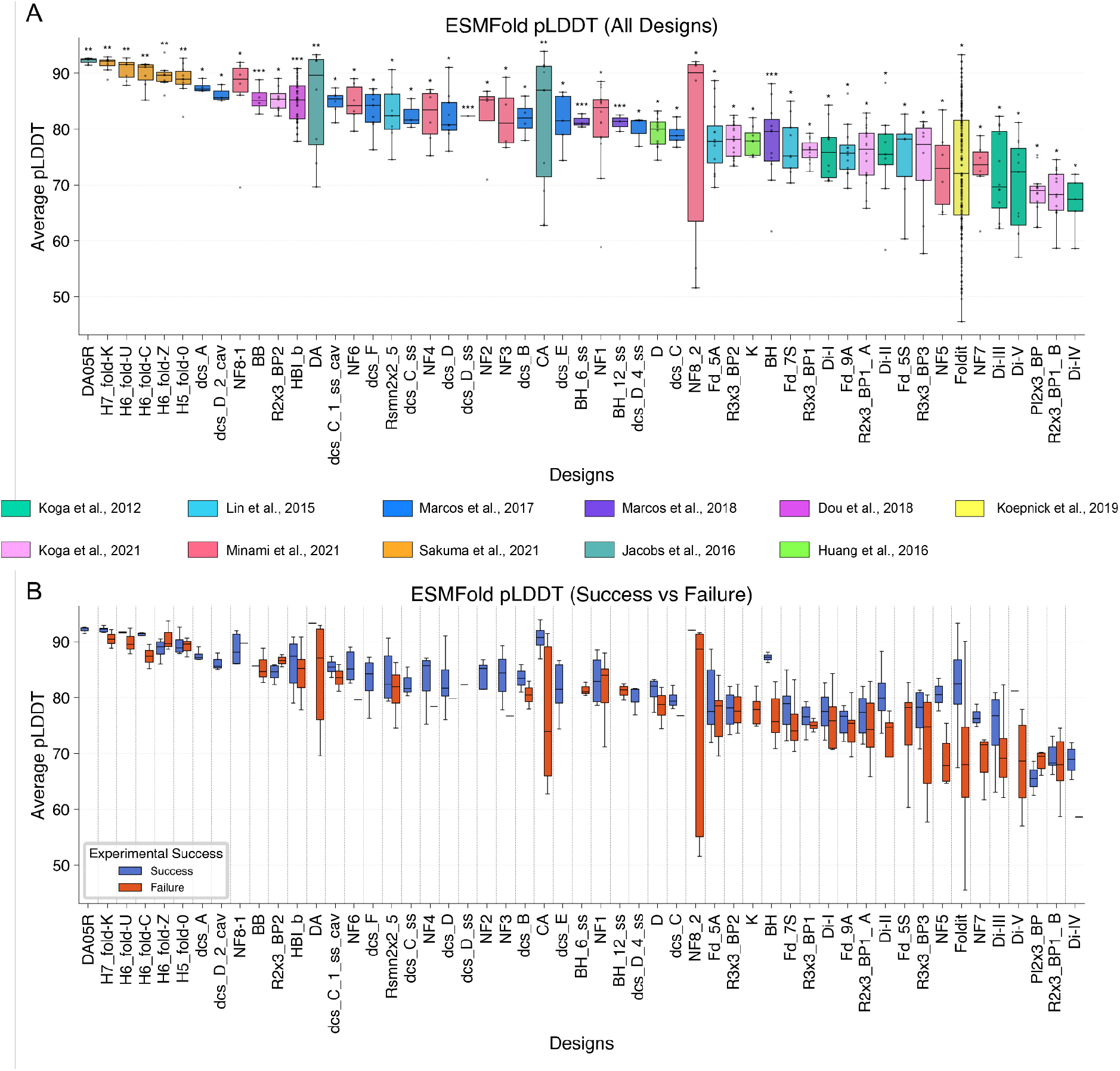
ESMFold average pLDDT distributions for designed topologies. **A)** Distribution of ESMFold average pLDDT for designs included in this study. **B)** Distribution of ESMFold average pLDDT for experimental successful designs (blue) and unsuccessful designs (orange) for each design topology

**Figure S6.**
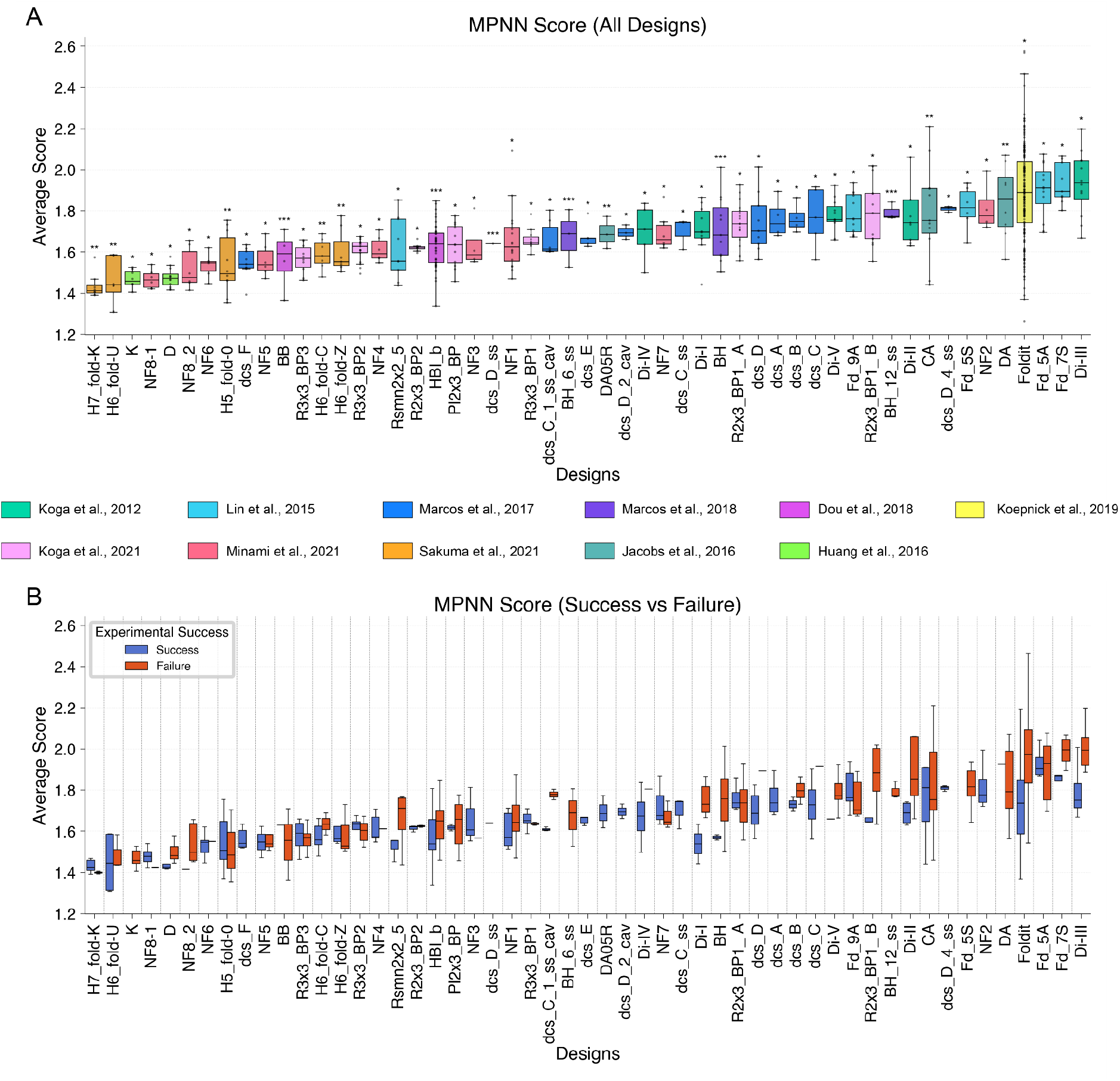
ProteinMPNN score distributions for designed topologies. **A)** Distribution of MPNN scores for designs included in this study. **B)** Distribution of MPNN score for experimental successful (blue) and unsuccessful designs (orange) for each design topology.

